# Different B cell activation patterns in asymptomatic and symptomatic COVID-19 patients

**DOI:** 10.1101/2022.12.19.521064

**Authors:** Nhung Pham, Nuray Talih, Friederike Ehrhart, Chris T Evelo, Martina Kutmon

**Affiliations:** Department of Bioinformatics - BiGCaT, School of Nutrition and Translational Research in Metabolism (NUTRIM), Maastricht University, Maastricht, Netherlands; Maastricht Centre for Systems Biology (MaCSBio), Maastricht University, Maastricht, Netherlands

**Keywords:** B cells, COVID-19, molecular pathway

## Abstract

Early and persistent defects in B cell subsets such as memory B cells were shown to be correlated with poor outcomes in COVID-19 patients. This research aimed to develop a molecular pathway model to understand the B cell development in COVID-19. A B cell transcriptomics dataset, obtained from COVID-19 patients, was analyzed on the resulting pathway model to study B cell activation. The pathway showed two distinct gene expression profiles between asymptomatic and symptomatic patients. In asymptomatic patients, there is an increase in transcript levels of antiviral interferon-stimulated genes such as ISG15, IFITM1, and NEAT1 and a driving gene for the extrafollicular pathway CXCR4 indicating a formation of plasmablast. In symptomatic patients, the results suggest an inhibition occurring at the germinal center hinting at a reduction in memory B cell production. Transcripts of driver gene CXCR5 involved in germinal center development is one of the most downregulated genes. This could contribute to the shortage in the formation of memory B cells in COVID-19. Concluding, in SARS-CoV-2 infection, B cells follow different activation routes in asymptomatic and symptomatic patients. In this study, we constructed a pathway that allowed us to analyze and interpret activation patterns of B cells in COVID-19 patients and their link to disease severity. Importantly, the pathway and approach can be reused for further research in COVID-19 or other diseases.

## 1. Introduction

Recent COVID-19 studies showed a correlation between delayed and weak adaptive immune responses with poor clinical outcomes in severe patients (1; 2). Part of adaptive immunity is the humoral response has been shown to be delayed in severe COVID-19 patients (2). In the humoral immune response, B cells are essential for presenting antigens to other immune cells, producing pathogen-specific antibodies, and secreting proinflammatory cytokines. Low B cell counts of some subsets such as memory B cells and expression of B cells related genes such as heavy chain and light V chain genes are observed in severe COVID-19 patients (3; 4; 5).

B cells originate from hematopoietic stem cells in the bone marrow, where they differentiate into immature B cells. These immature B cells then migrate via the blood to secondary lymphoid organs, such as the spleen or the lymph nodes, to complete maturation. Upon infection or immunization as a response to an invading pathogen, the circulating mature but naïve B cells are activated through antigen encounter and may follow one of two pathways to differentiate into antibody-secreting cells. They may either develop along canonical responses involving germinal center reaction, where they can differentiate into long-lived B cell populations, namely memory B cells, and plasma cells (6; 7; 1). Alternatively, B cells may participate in non-canonical responses that lack GCs and cooperate in B cell proliferation and differentiation into short-lived plasmablasts at extrafollicular sites (8). Both pathways ultimately lead to the production and secretion of antibodies in the form of antibody-secreting cells (6; 7; 9).

In this study, we aim to create a molecular pathway model of B cell activation in COVID-19 to comprehend the transformation of B cells into different subsets through different differentiation routes. The model will then be used for data analysis to identify key regulators of the pathway activation in COVID-19 in correlation to disease severity.

## 2. Material and Methods

### 2.1. Pathway construction

The pathway model was constructed following standard guidelines (10). A literature review was done initially, to collect all relevant information. Besides studies on B cell activation in relation to SARS-CoV-2, generalized research on B cell activation in response to infection was also used. As a starting point, the pathway was built based on a diagram that depicts the migration of B cells from the outer lymph node into the follicle upon activation from Cerutti et al (11). The pathway model was then constructed using an open-source pathway analysis and drawing software PathVisio version 3.3.0 (12). Gene products and proteins were annotated with identifiers from Ensembl (13) and UniProt (14). Literature references were added to specific sections and interactions in the pathway. The pathway model was published on WikiPathways, a community database of curated biological pathways (15).

### 2.2. Pathway visualization

A single-cell transcriptomic dataset obtained from B cells of 130 patients infected with SARS-CoV-2 with varying severity (asymptomatic, mild, moderate, severe, and critical) (16) was used to study the activation of B cells in the curated pathway model. The dataset included RNAseq data from three different B cell types: immature, naive, and exhausted B cells. Only data for naive and exhausted B cells were used in this study because our pathway model focuses on the activation of B cells after the immature stage. Differentially expressed genes were identified as those with an adjusted p-value *<* 0.05. Expression values of these genes were then projected and visualized on the pathway model using Cytoscape version 3.9.1 (17) and the WikiPathways app version 3.3.7 (18).

## 3. Results

### 3.1. B cell activation in COVID-19: pathway description

To study the B cell development during SARS-CoV-2 infection, we constructed a machine-readable molecular pathway model based on 25 papers of which five are COVID-19 specific. The final pathway model depicts the initial B cell activation inside the lymph node and the following activation routes to differentiate to either plasmablast cells following the extrafollicular pathway or memory B cells through the follicular pathway (Figure 1). The pathway was deposited in WikiPathways under the identifier WP5218 (19).

**Figure 1:**
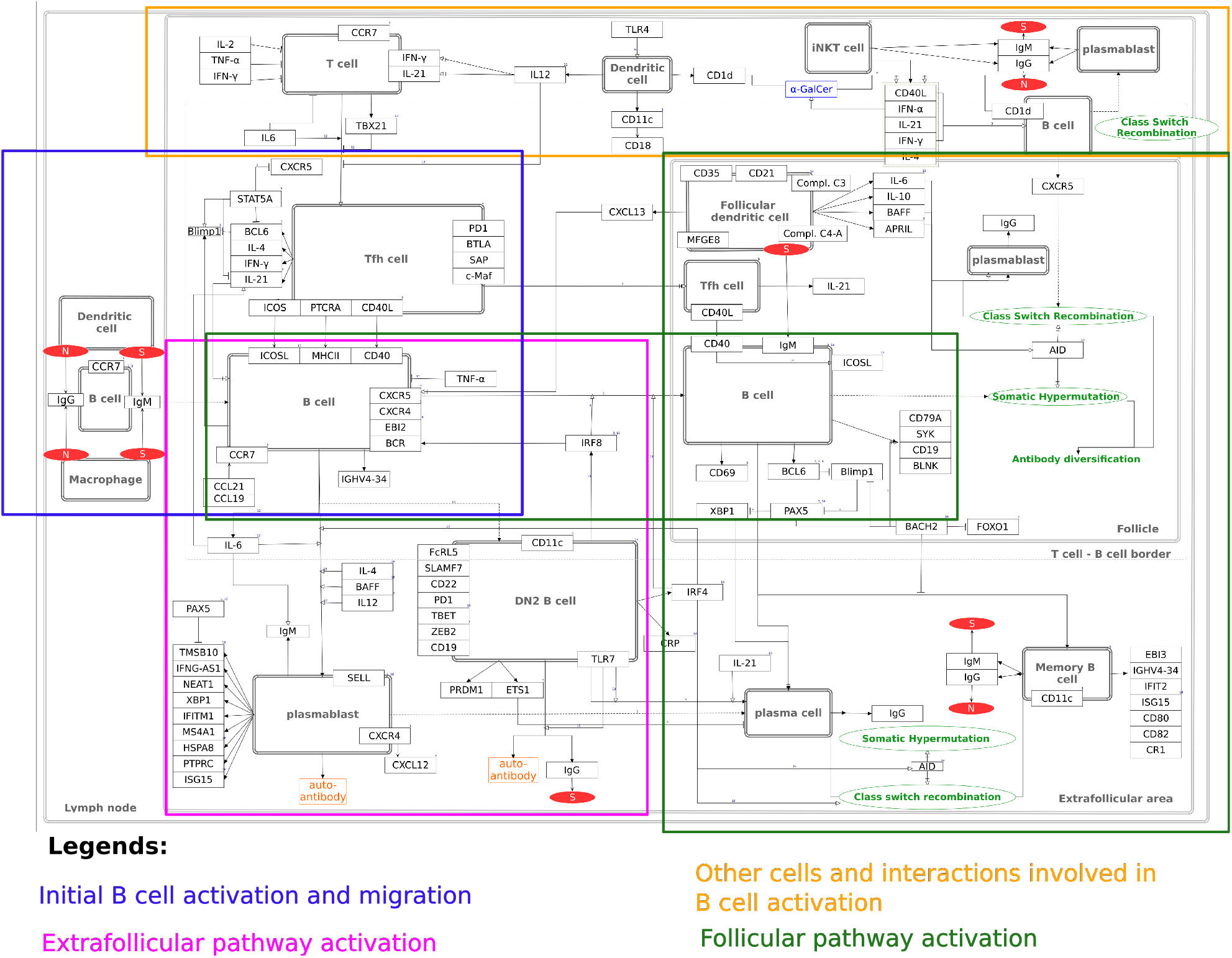
Extrafollicular B cell activation by SARS-CoV-2 pathway. The boxes indicate different processes that occur during B cell development. The full-size resolution figure can be found on www.wikipathways.org/instance/WP5218.

#### 3.1.1. Initial B cell activation and migration (Figure 1, blue box)

Initially, in the lymph node, naive B cells can interact with either macrophages or dendritic cells that carry SARS-CoV-2 antigens (N and S proteins) on their surface (20; 21). Via its IgM and IgG receptors, the B cell receives activation signals to process the antigen and subsequently migrate to the T-B border of the lymph node (11; 22; 23; 24). A balanced expression of CCR7 and CXCR5 mediates B cells to migrate to the interface between follicles and T-cell areas (T-B border zone) (25; 26; 27). At the T-B border, the B cell displays a peptide MHC-II antigenic complex, through which it can interact with the pre-T-cell antigen receptor alpha of an early state T follicular helper (TFH) cell. Through its CD40L receptor, the TFH cell can bind to the CD40 receptor of the B cell. Furthermore, the TFH cell secretes the cytokines IL21, IL4, and IFN*γ*, all of which stimulate the development of the B cell (11). When the B cell gets activated, EBI2 expression is greatly increased. EBI2 is necessary for the localization of the B cell (27). Downregulation of EBI2 indicates GC differentiation whereas the persistence of the EBI2 receptor suggests the integration of the activated B cell into the extrafollicular pathway (28; 29). Apart from this, the factors IRF4 and IRF8 are also involved in B cell guidance. High concentrations of IRF4 as a consequence of B cell activation drive the rapid generation of plasmablasts (30). Besides facilitating antigen-specific interaction with T helper cells, IRF8 in contrast promotes GC differentiation (31). Following the interaction with the TFH cell, the B cell may enter either the canonical (follicular) or extrafollicular pathway.

#### 3.1.2. Canonical/ follicular pathway activation (Figure 1, green box)

For the B cell to follow the canonical pathway and travel inside the follicle, it has to upregulate the expression of CXCR5 (32). This is stimulated by the chemokine CXCL13, which also excites the movement of TFH cells toward the follicle (32). CXCL13 itself is expressed by follicular dendritic cells (FDCs) (33). FDCs are a major reservoir for antigens and play a crucial role in the formation of germinal centers (GCs) (34). The FDC also expresses BAFF (35; 34; 36), IL6 (9), IL10 (36), and APRIL (11; 36), all of which modulate B cell differentiation within the GC. In the second step, the CD40 receptor of the B cell will undergo a cognate interaction with the CD40L receptor of the TFH cell (37). Furthermore, the CD40 receptor can stimulate the expression of ICOSL, which then promotes the formation of plasma cells (38; 11). Consequently, the B cell may undergo Class Switch Recombination and affinity maturation through Somatic Hypermutation, ultimately leading to the diversification of antibodies (11). Upon encounter with their cognate antigen, B cells rapidly upregulate CD69 expression (39). Amongst others, GC B cells can be recognized by their expression of BCL6, which inhibits the differentiation towards plasmablasts by repressing BLIMP1 (40). BLIMP1 in turn suppresses PAX5, which acts as a dual function activator and suppressor and is essential for GC formation (40). For instance, PAX5 represses XBP1, involved in plasma cell differentiation (41). Along with BACH2 and BCL6, PAX5 stabilizes the GC B cell state, while antagonizing plasma cell differentiation. Following GC formation, B cells may either develop into long-lived memory B cells or into plasma cells that primarily produce IgG (11), depicted in the lower corner on the right side of the pathway model. In case the B cell differentiates into an IgM or IgG antibody producing memory B cell, the factors EBI3 (42), IGHV4-34, IFIT2, ISG15, CD80, CD82, and CR1 will be expressed (43).

#### 3.1.3. Extrafollicular pathway activation (Figure 1, pink box)

While CXCR5 is an indicator of B cell maturation towards the GC, an increased expression of CXCR4 may in turn portend toward an extrafollicular pathway entry (26). Differentiation of the B cell following the non-canonical pathway ultimately leads to the generation of short-lived plasmablasts that secrete IgM. This process is stimulated by IL4, BAFF, IL12, and IL6, which are produced by innate immune cells and all promote B cell activation and maturation (11). Hence, dysregulation of B cells may therefore be caused by the innate immune response. Downregulation of CXCR5 and CCR7 then direct plasmablast migration towards extrafollicular growth by decreasing their responsiveness to the follicular and T zone chemokines CXCL13, CCL19, and CCL21 (44). The resulting plasmablast populations express the PAX5 repressed gene TMSB10 (31; 45), as well as the genes XBP1 (46), NEAT1, MS4A1, HSPA8, PTPRC, IFITM1, ISG15, and IFNG-AS1, that indicate an ongoing interferon response (43). Particularly, ISG15-secreting plasmablasts are a proinflammatory feature in autoimmune diseases like SLE (47). Plasmablasts can differentiate into plasma cells via the association of CD11c high dendritic cells (48).

Alternatively, to directly develop into a plasmablast via the extrafollicular pathway, B cells may also develop into double negative type 2 (DN2) B cells, which are closely related to activated naïve B cells and are prone to differentiate into plasma cells. DN2 B cells may undergo class switching, however, they lack IgD and CD27 expression. They are a unique type of memory B cells that are expanded in a variety of diseases, particularly autoimmune diseases, and are known to contribute to pathogenesis (49). The formation of DN2 B cells is indicated by several markers, including FCRL5, SLAMF7, CD22, PD1, TBET, ZEB2, and CD19 (49; 36). Furthermore, DN2 B cells express low amounts of ETS1 gene which may inhibit plasma cell differentiation (36). When ETS1 is deficient, this may cause an extrafollicular accumulation of autoreactive plasma cells. On the other hand, TLR7 and IL21 stimulate DN2 B cell maturation into plasma cells (30). Besides, TLR7 may also enhance IgG antibody production but is also involved in causing the production of autoantibodies (30).

#### 3.1.4. Other cells and interactions involved in B cell activation (Figure 1, yellow box)

Although the main focus of this pathway is the B cell development into antibody secreting cells, other cells also play an important role in guiding the B cell toward maturation. For instance, before the T cell is equipped with B cell helper activity, it needs to be activated by a dendritic cell. The early T cell resides in the extrafollicular area of the lymph node, depicted in the upper left corner of the model. The dendritic cell itself expresses high amounts of CD11c and is stimulated by TLR4 (48). Regulation of humoral immunity toward short-lived extrafollicular responses occurs via the expression of IL12, which is known to inhibit TFH differentiation strongly (9). Other factors inhibiting TFH cell development include IFN*γ*, Il21, IL2, and TNF*α* (9). While TBX21 suppresses GC responses, IL6 indirectly promotes GC formation through the stimulation of TFH differentiation (9). The activated T cell may further migrate to the T-B border.

The top right corner of the pathway resembles the interaction between an invariant natural killer T cell (iNKT cell) with a B cell. When the B cell presents its glycoprotein CD1d, the iNKT cell gets stimulated to express CD40L, IFN*α*, IFN*γ*, IL4, and IL21, which enables further differentiation of the iNKT cell and aids the differentiation of B cells into extrafollicular plasmablasts that secrete IgG and IgM (11). The iNKT cell also promotes the cognate interaction of the dendritic cell with the T cell, which may then differentiate into a TFH cell. This occurs via the presentation of *α*-galactosylceramide (*α*-GalCer) of the dendritic cell to the iNKT cell, promoting the interaction of the dendritic cell with a T cell (11).

### 3.2. B cell development pattern in COVID-19 patients with various degrees of severity

The resulting pathway model was used to visualize the public transcriptomics data of two different B cell types: naive and exhausted B cells obtained from patients with five degrees of severity: asymptomatic, mild, moderate, severe, and critical (16).

All 73 genes in the pathway were found in the dataset. Among them, 33 genes (59 %) are significantly influenced by SARS-CoV 2 with an adjusted p-value smaller than 0.05 (Figure 2). 16 genes are upregulated and 21 genes are downregulated in SARS-CoV 2 infection in at least one cell type and one degree of severity.

**Figure 2:**
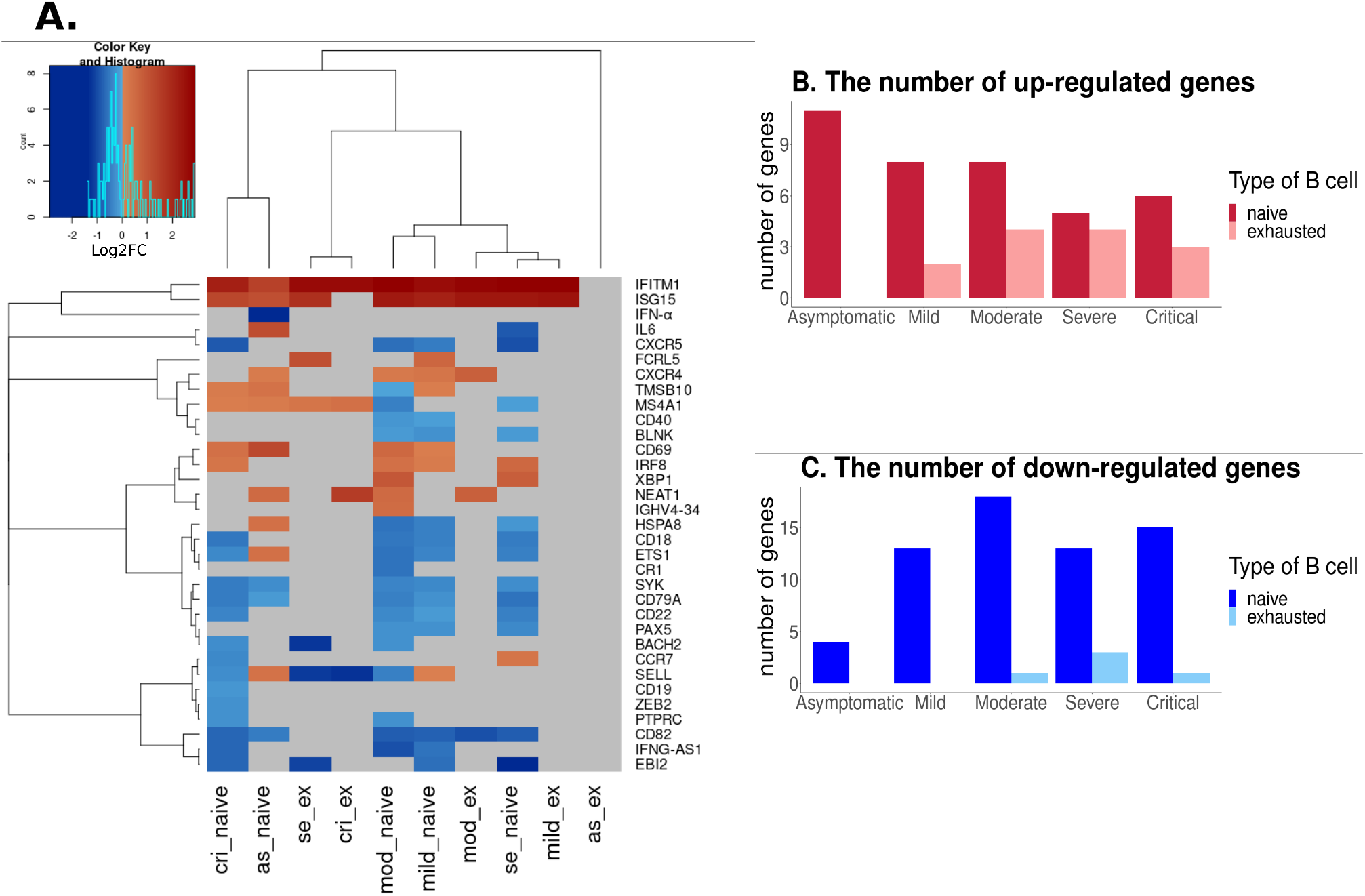
Gene expression in naive and exhausted B cells in patients with various degrees of severity. **A**. Heatmap of gene expression in different B cell types in patients with various degrees of severity; **B-C**. The number of up- and downregulated genes in varying SARS-CoV-2 infection severity and proceeding B cell development. Red-upregulated genes, blue - down regulated genes, grey-not measured. As : asymptomatic patients; cri : critical patients; se : severe patients; mild :mild patients; mod : moderate patients; im: immature B cell; naive: naive B cell; ex: exhausted B cell.

The pathway showed a clear distinct gene expression profile between asymptomatic and symptomatic patients. Among patients, the pathway showed the most upregulated genes and the least downregulated genes in asymptomatic patients (Figure 3). In these patients, B cells tend to go toward short-lived plasmablast cells with a high expression of antiviral interferon-stimulated genes such as ISG15, IFITM1, and NEAT1.

**Figure 3:**
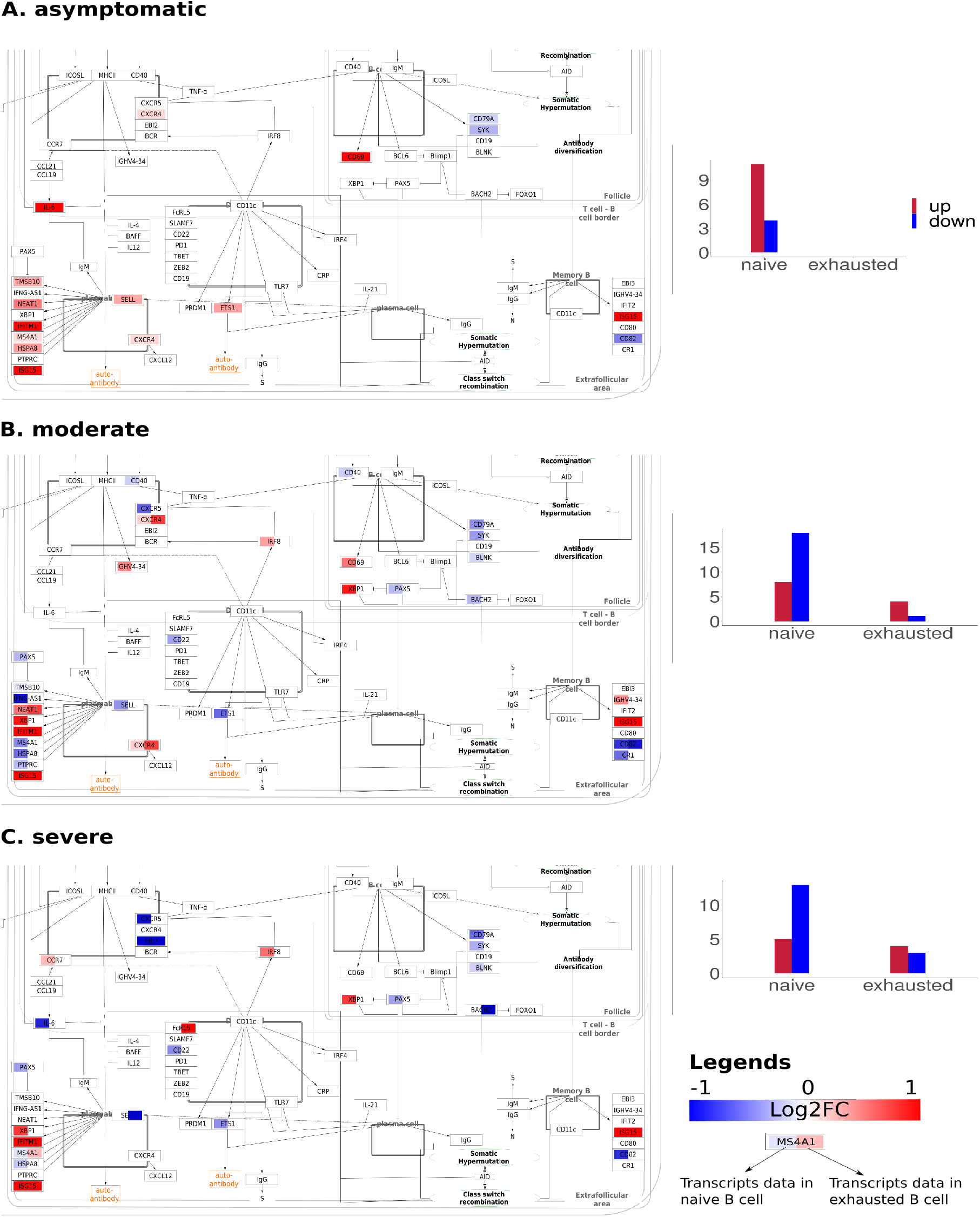
Data visualization in asymptomatic, moderate and severe patient. The pathway showed a clear distinction in gene expression between asymptomatic and symptomatic patients. **A**. In asymptomatic patients there are more upregulated genes (red) than downregulated genes (blue); **B-C**. In symptomatic patients, there are more downregulated genes. Each gene in each pathway is visualized with data from naive (left box) and exhausted B cells (right box). In asymptomatic patients, only data from naive B cells are significant for visualization. Visualization in mild and critical patients are provided in the supplementary file 1.

In asymptomatic patients, the pathway is affected the least with the least total number of genes changed in expression compared to patients with more severe symptoms. The pathway showed the most genes affected in moderate patients (Figure 2BC). In severe patients, the gene has the biggest changes in their transcriptome level (Figure 2A). In these patients, almost all B cell types show down-regulation of marker genes such as SELL, CD22, and CD19.

Most genes driving B cells toward the follicular area such as CXCR5 are down-regulated in symptomatic patients with more reduction when increasing severity. Essential genes for the stabilization of GC formation such as PAC5 and BACH2 are slightly downregulated in almost all conditions. Furthermore, most of the genes expressed by the follicular B cell such as CD79A, SYK, CD19, and BLNK appear rather downregulated. On the contrary, CXCR4, which supports extrafollicular development, occurs to be slightly upregulated only in asymptomatic, mild, and moderate patients. Across cell types, naive B cells have the most genes influenced in the pathway, especially in critical and moderate patients. During B cell maturation, the dysregulation of genes decreases for the exhausted B cell state.

Among all changed transcript levels in the pathway, IFITM1, ISG15, CD69, and FCRL5 are the most upregulated ones (Figure 2A). IFITM1 is upregulated in almost all cell types and severity levels except in exhausted B cells in asymptomatic patients. The highest expression of IFITM1 was found in severe and moderate conditions and is lowest in asymptomatic patients. ISG15 is upregulated in both naive and exhausted B cells and severity levels except for exhausted B cells in asymptomatic and critical patients. ISG15 transcript is highest in severe patients and the lowest in asymptomatic patients. CD69 is upregulated in naive B cells of all patients except for severe patients. FCRL5 is upregulated in naive B cells of mild patients and exhausted B cells of severe patients. The most downregulated transcripts in the pathway are IFN*α*, EBI2, BACH2, and CXCR5. IFN*α* is downregulated in naive B cells of asymptomatic patients. EBI2 is downregulated in naive B cells of mild, severe, and critical patients and in exhausted B cells of severe patients. BACH2 is downregulated in naive B cells of moderate and critical patients, and in exhausted B cells of severe patients. CXCR5 is downregulated in naive B cells of all patients except for asymptomatic patients. The transcript level of CXCR5 is the most downregulated in severe and critical patients.

Some transcripts have opposite expression levels across cell types and patients. SELL is upregulated in naive B cells of asymptomatic and mild patients but downregulated in naive B cells of moderate and critical patients and in exhausted B cells of severe and critical patients. IL6 is upregulated in naive B cells of asymptomatic patients but downregulated in the same cell of severe patients.

## 4. Discussion

To understand the B cell activation in COVID-19 patients, we constructed a molecular pathway model. The pathway depicts B cell differentiation into short-lived plasmablast cells in the extrafollicular area and memory B cells or long-lived plasma cells in the follicular area. We further analyze the behaviors of B cells in COVID-19 patients using published transcriptomics data.

In our analysis, we found an increase in the expression of ISG15, CXCR4 a driving gene for the extrafollicular pathway and an early-antigens detector marker CD69 in asymptomatic patients. This upregulation could indicate that asymptomatic patients quickly recognized and responded to the virus with the help of plasmablast cells in the extrafollicular pathway. A recent study by Hinai et al. (2022) (50) has shown that an increase in the expression of interferon stimulated genes reduces the viral copies of influenza A virus in vitro. The upregulation in interferon stimulated genes in asymptomatic COVID-19 patients was also found in other studies and was correlated with good clinical outcomes in these patients (51; 52). The upregulation in the formation of plasmablast and no change in essential genes for germinal center formation in asymptomatic patients from our study support the hypothesis that asymptomatic patients have a quick and effective innate immune response to control viral replication which helps to prevent a severe clinical outcome (53; 54).

In symptomatic patients, the data visualized in the pathway shows upregulation of genes in the extrafollicular area indicating plasmablast formation but also downregulation in genes in the follicular area indicating a lack of canonical pathway activation or germinal center activation. Especially, the more severe the disease the more reduction in gene expression in the follicular area. These findings are consistent with previous findings in literature (55; 56; 57; 16). We furthermore found that CXCR5, a key regulator of follicular pathway activation, indicating follicle entry of B cells, is downregulated in all cell types and severity levels in symptomatic patients, especially in severe and critical cases. This indicates that the B cell maturation in these patients is guided towards the extrafollicular pathway. B cells lacking CXCR5 have been shown to be correlated with disease severity and poorer outcomes in human systemic lupus erythematosus (58). A study from Shuwa H.A. et all (2021) reported that B cells from acute COVID-19 patients display a significantly reduced expression of chemokine receptors CXCR3 and CXCR5 (59).

### 4.1. Limitation and next steps

The current study has some limitations. B cell development and differentiation process is a complex process with many mediate steps. In order to maintain the interpretation of the pathway, many intermediate B cell types and gene products are not included in the model. In addition, most of the literature used for constructing the pathway is from general or immune responses to other diseases due to the lack of specific data on B cells in COVID-19. Nevertheless, WikiPathways is an open platform, the pathway can serve as a foundation for further curation and new knowledge on COVID-19 can be easily added to the pathway.

## 5. Conclusion

By analyzing the published transcriptomics dataset on the pathway molecular model, we identified two distinct activation patterns of B cells in asymptomatic and symptomatic COVID-19 patients. We confirmed key findings that have been described in the literature and demonstrated a new, intuitive way to visualize and analyze omics data. We demonstrated the usefulness of a molecular pathway model for helping to interpret biological outcomes from omics data. With the current pathway model, the same research could be done quickly for other diseases and for the comparison of different disease signatures.

## Data availability statement

The pathway model constructed from this study can be found at www.wikipathways.org/instance/WP5218.

## Ethics approval statement: human and animal studies

Not relevant

## Patient consent statement

Not relevant

## Permission to reproduce material from other sources

Not relevant

## Clinical trial registration

Not relevant

## Author contributions

NP, NT and MK conceived the idea. NT and NP constructed the pathway. NP conducted the data analysis. NP interpreted the result. NP and NT draft the manuscript. MK, FE, and CE edited the manuscript. All authors contributed to the article and approved the submitted version.

