## Supplementary figure 1 for "Different B cell activation patterns in asymptomatic and symptomatic COVID-19 patients"

### Supplementary file

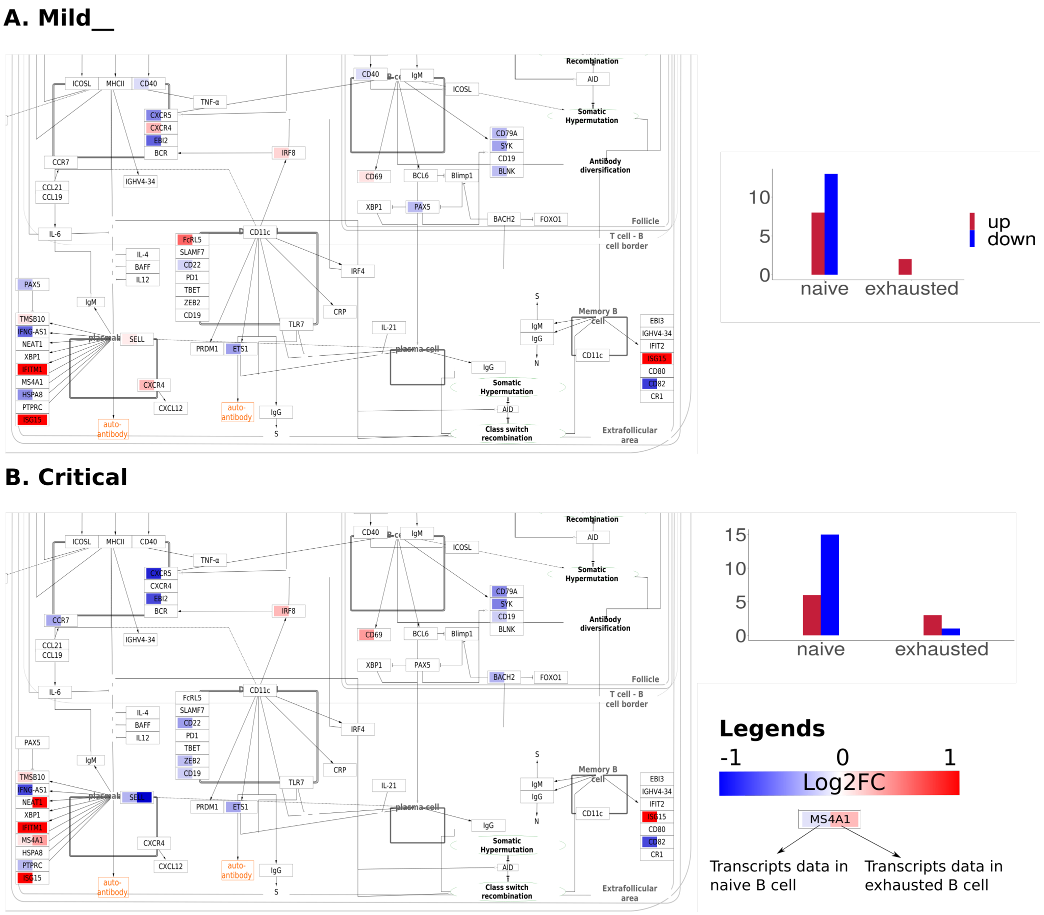


Figure 1: **Data visualization in mild and critical patients**. In these patients, there are more downregulated genes with stronger reduction when increasing severity. Each gene in the pathway is visualized with data from naive (left box) and exhausted B cells (right box).
